# Morphodynamic Atlas for Drosophila Development

**DOI:** 10.1101/2022.05.26.493584

**Authors:** Noah P Mitchell, Matthew F Lefebvre, Vishank Jain-Sharma, Nikolas Claussen, Marion K Raich, Hannah J Gustafson, Andreas R Bausch, Sebastian J Streichan

## Abstract

During morphogenesis, diverse cell-scale and tissue-scale processes couple to dynamically sculpt organs. In this coupling, genetic expression patterns and biochemical signals regulate and respond to mechanical deformations to ensure reproducible and robust changes in tissue geometry. A long-standing approach to characterize these interactions has been the construction of expression atlases, and these atlases have necessarily relied on fixed snapshots of embryogenesis. Addressing how expression profiles relate to tissue dynamics, however, requires a scheme for spatiotemporal registration across different classes of data that incorporates both live samples and fixed datasets. Here, we construct a morphodynamic atlas that unifies fixed and live datasets – from gene expression profiles to cytoskeletal components – into a single, morphological consensus timeline. This resource and our computational approach to global alignment facilitate hypothesis testing using quantitative comparison of data both within and across ensembles, with resolution in both space and time to relate genes to tissue rearrangement, cell behaviors, and out-of-plane motion. Examination of embryo kinematics reveals stages in which tissue flow patterns are quasi-stationary, arranged as a sequence of ‘morphodynamic modules’. Temperature perturbations tune the duration of one such module – during body axis elongation – according to a simple, parameter-free scaling in which the total integrated tissue deformation is achieved at a temperature-dependent rate. By extending our approach to visceral organ formation during later stages of embryogenesis, we highlight how morphodynamic atlases can incorporate complex shapes deforming in 3D. In this context, morphodynamic modules are reflected in some, but not all, measures of tissue motion. Our approach and the resulting atlas opens up the ability to quantitatively test hypotheses with resolution in both space and time, relating genes to tissue rearrangement, cell behaviors, and organ motion.

## I. INTRODUCTION

There is a long history of constructing transcriptomic and protein expression atlases to tackle outstanding questions in morphogenesis [1–7]. Existing atlases have been, by necessity, static representations of embryo components at a collection of specific points in time [4, 8, 9]. These bottom-up approaches have proven useful in elucidating otherwise-hidden connections between disparate components of the morphogenetic program. However, morphogenesis consists of integrated dynamic processes, subject to dynamic rules. Therefore, a full understanding of how biological shape emerges requires a synthesis of spatial and temporal information. This demands new methods for registering independent datasets in both space and time.

Not all proteins of interest can be visualized in the same recording, and constructs for live imaging are often unavailable. Therefore, a morphodynamic atlas has to combine data from different experiments, and from fixed as well as from live samples. This poses three challenges: 1) spatial alignment to compare different features of a given tissue region imaged in different experiments, (2) temporal alignment to compare how features from different experiments relate over time, and 3) timeline construction, creating a single morphogenetic timeline applying to all experiments. To create an efficient and rigorous scientific resource, all of these steps need to be automated and carried out quantitatively.

Here we provide a solution to this challenge by generating a dynamic protein expression atlas that integrates datasets spanning much of Drosophila embryogenesis. We demonstrate the analytical value of our atlas by first focusing on the earliest movements of gastrulation. We then generalize to visceral organ morphogenesis, demonstrating that our approach to spatiotemporal registration of dynamic datasets applies also to more complicated shapes deforming in 3D.

Our atlas consists of over 500 unique fixed and live embryo datasets, including 18 mutant genotypes, detailed in the Supplementary Information. All data was captured using *in-toto* multi-view light sheet microscopy – enabling global analysis of dynamics [10–12]. The atlas defines temporal coordinates based on tissue morphology and spatial coordinates based on tissue cartography. We generated a fully documented, MATLAB-based platform for querying data classes of choice, including expression profiles, tissue flows, and expression anisotropy [13]. The atlas platform is an open-source tool specifically designed to incorporate future data contributions.

## II. CONSTRUCTION OF A MORPHODYNAMIC ATLAS OF EARLY DROSOPHILA GASTRULATION

We integrate independent classes of experimental data onto a fixed two dimensional frame of reference indexed along a single morphological timeline. Fig 1 provides a visual example of this approach. Both live and fixed expression data — such as gene products involved in patterning (pair-rule genes (PRG)) and cytoskeletal components (myosin and integral junction proteins) — are visualized using an iterative cartographic approach which makes 2D projections of the embryo’s 3D surface (Fig. 1). By imaging the samples *in toto,* we can use the full embryo geometry to perform spatial alignment. Our software automatically aligns a cartographic projection of the 3D embryo surface with the embryo body axes for this purpose (see Methods). This method enables rapid calculation and comparisons of tissue dynamics and expression patterns across embryos, with each dataset providing expression patterns at subcellular resolution.

**FIG. 1.**
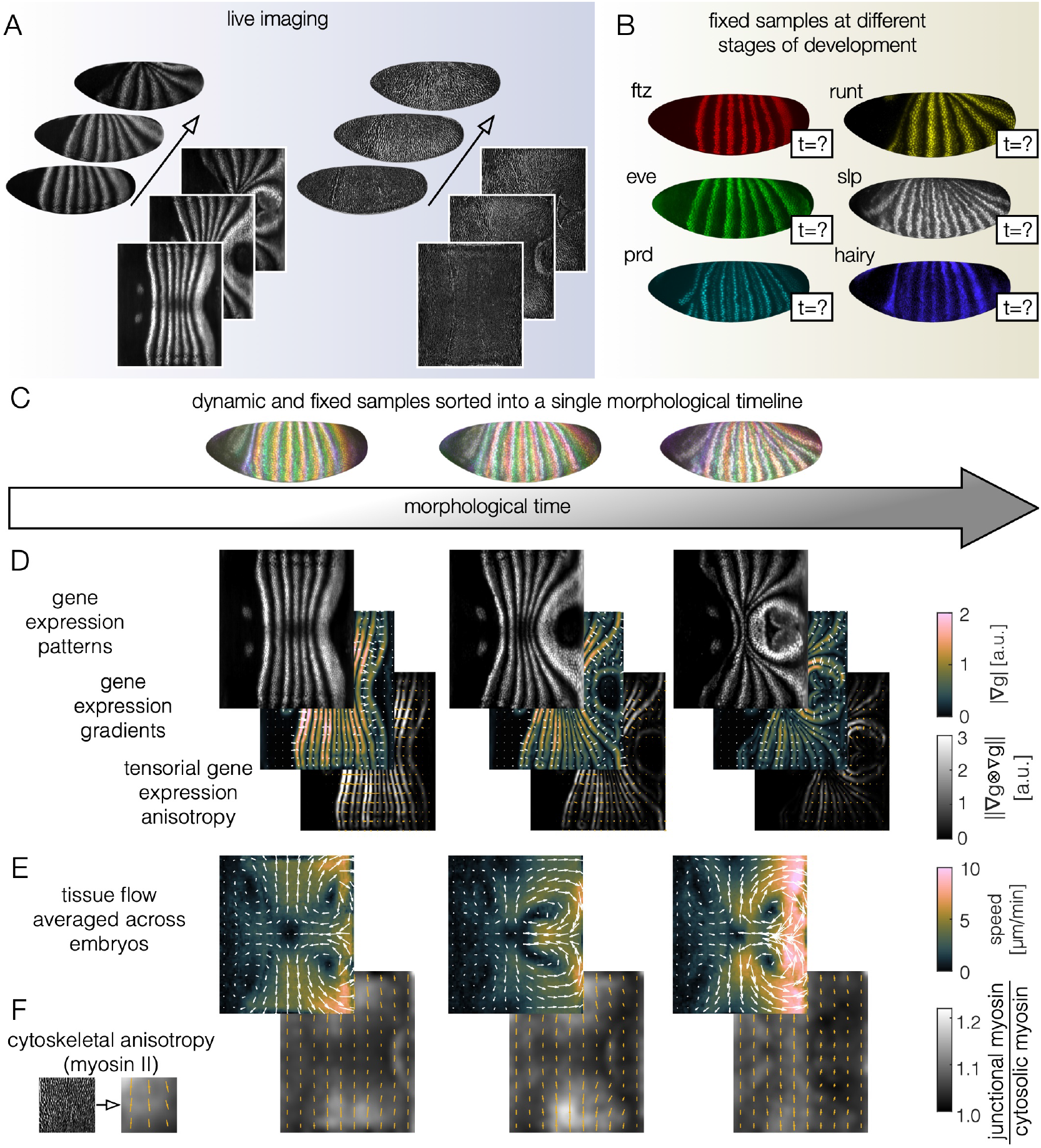
A single morphological timeline parameterizes different classes of dynamic and fixed data during Drosophila axis elongation. *(A)* Confocal lightsheet imaging of hundreds of embryos generates volumetric datasets, which we spatially register into a fixed 2D parameterization. Here we show representative snapshots of a Runt nanobody reporter and myosin marker, sqhGFP. *(B)* Fixed embryos show spatial patterning, but require a method for generating timestamps. *(C-F)* Registering all data to a global morphological timeline enables quantitative characterization of scalar fields – such as transcription factor expression levels (top row), vector fields – such as gradients of expression or tissue velocities, and tensor fields – such as measures of cytoskeletal anisotropy, or tensors computed from gradients of gene expression. Panel (E) shows average velocity fields across five morphologically aligned sqh:GFP datasets. Panel (F) shows the tensorial myosin patterning via the radon transform across the same sample (c.f. [17]), and the colorbar represents the norm of the deviatoric (anisotropic) component of the myosin tensor.

A feature of this technique is the capacity to describe the dynamic morphogenetic program in terms of underlying physical *fields*. The position of individual cells will vary from sample to sample, so single-cell comparison across embryos is not possible. Instead, we extract mesoscale information by smoothing gene expression patterns, velocity fields, and measures of anisotropy. These fields can then be related to one another across the embryo’s body plan, for example, allowing us to look for large-scale correlations of patterns.

Recently, this biophysical field-theoretic approach has led to fruitful models relating mechanical force, gene expression, and tissue geometry [14–19]. Fig. 1C highlights a subset of the ways in which a single dataset can be described using multiple distinct classes of spatiotemporal fields. Our atlas therefore provides ready access to spatially-aligned fields, ripe for quantitative hypothesis testing.

While our cartographic pipeline solves the problem of aligning different recordings in space, these recordings also have to be aligned in time. Within a population of embryos, morphogenesis proceeds at different rates, due to natural variation between individuals that is exacerbated by genetic factors and ambient environmental conditions. This raises two central questions: what is the relevant timeline for analysis of morphogenetic processes, and how do we consolidate disparate timelines? For centuries, the standard technique used to track developmental progress has been to define stages based on the appearance of canonical morphological milestones. Here, we adopt this paradigm of timing morphogenesis using tissue geometry. However, rather than only timing discrete stages, we upgrade the characterization of morphological time into a continuous variable suitable for quantitative analysis.

We proceed in two steps: based on pairwise comparison of suitable morphological features, (1) the timelines of individual live datasets are dilated to align along a consensus timeline, and (2) fixed samples are placed appropriately in the consensus timeline.

To carry out step (1), we first select a suitable morphological feature which allows us to score the similarity of two images (see Sect. II A for a concrete example). To align two live datasets, one could simply find the fixed delay between the two which maximizes average similarity (e.g. concluding that movie A begins 10 minutes before movie B). However, a rigid alignment is incapable of taking into account the variable rates at which morphogenesis proceeds in different embryos. We instead compare every frame of one movie to every frame of the other, creating a rectangular matrix of correlations (Fig. 2). A fastmarching algorithm then finds the optimal path through this matrix, matching every timepoint of movie A to its best match in movie B while respecting temporal order, naturally accounting for variable rates of morphogenesis in the two samples.

**FIG. 2.**
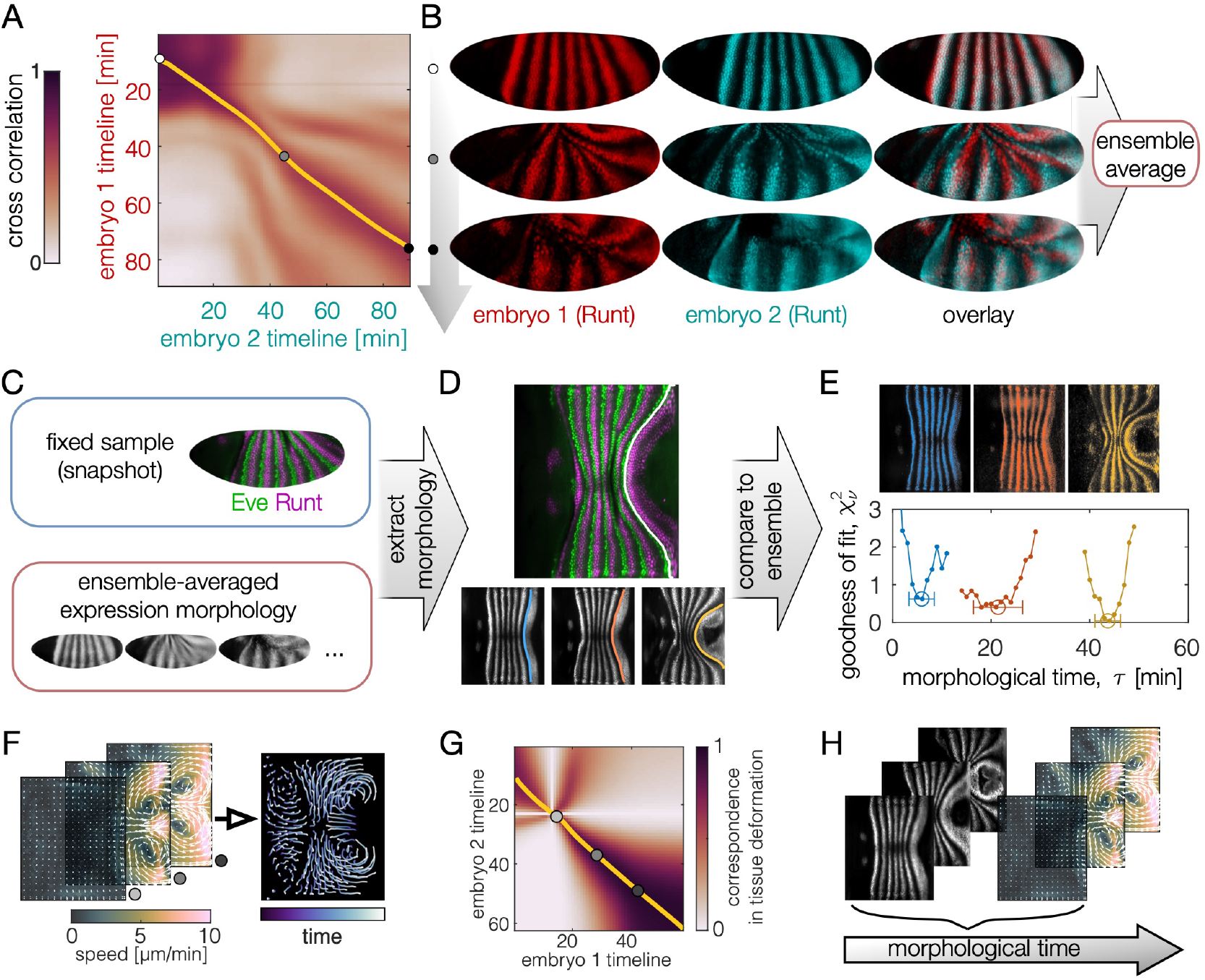
Live imaging of gene expresssion datasets define a consensus morphological timeline against which we timestamp fixed samples. *(A)* To build a common morphological timeline that can be used to timestamp fixed datasets, we compare each pair of live Runt nanobody datasets to one another, creating a matrix of frame-to-frame similarity values, through which we trace a correspondence curve (yellow). Annealing an ensemble of pairwise correlations such as this one yields a consensus timeline for ensemble averaging. For our consensus timeline based on live Runt nanobody data, the r.m.s. time difference between different correspondence curves was 2.1 minutes. (B) With pairwise correspondences in hand, we build a consensus timeline and overlay different embryos to create an ensemble average. (C) Fixed samples (top) are co-stained for Runt, allowing their comparison to the ensemble average of Runt nanobody live data. *(D)* We computationally extract the shape of Runt stripes of both the fixed sample (top) and the live data (bottom), allowing *(E)* for a quantitative comparison with the consensus timeline. For each fixed sample (here, we show three examples), this comparison measures the similarity to each point in the consensus timeline, yielding both a timestamp (circles) and an estimate of uncertainty (lateral errorbars). (F) For live datasets without Runt stripe expression, we integrate tissue velocities measured via PIV to record displacements. *(G)* Comparing tissue displacements across datasets (see Methods) leads to correspondence curves between live datasets. *(H)*Comparing integrated flows from live non-Runt datasets to those of live Runt datasets unites both in the morphological timeline.

### A. Aligning live datasets of PRG expression

As shown in Fig. 2, the PRG Runt is expressed in stripes along the embryonic dorsoventral axis that deform as the epithelium flows. By imaging a collection of embryos expressing a llama tag for Runt [20], we extract the continuously-deforming morphology of Runt stripes as a landmark for morphological timing to carry out steps (1) and (2) above. We thereby leveraged live datasets of Runt expression to build a consensus morphology of Runt stripe geometry. By linking the timelines for each pair of Runt embryos using our fast marching algorithm (see Methods), we generate a single, consensus timeline.

### B. Aligning fixed datasets

As shown in Fig. 2B, we designed our atlas such that all fixed embryos are multiply immunostained against both Runt and a second gene product target of interest. For each embryo, we extract the static geometry of Runt stripes to correlate its position against the consensus Runt stripe shapes from live datasets, thereby determining the timestamp at which the fit residual is minimum. This process uses stripe geometry as a stopwatch against which we timestamp all fixed samples in the atlas. Any co-stained targets of interest are therefore aligned in time based on the position of the Runt stripe.

While the capacity to align live and fixed datasets based on a shared protein expression feature is valuable, Runt expression data is not present in all live imaging datasets. In order to integrate these other types of live data, we therefore needed to extend our morphological approach.

### C. Aligning live datasets based on tissue deformation

Embryonic tissue deforms during morphogenesis, and its instantaneous rate of deformation can be captured by flow fields [17] (Fig. 3A). The degree of total deformation can serve as a benchmark for defining morphological time, and can be reconstructed from the instantaneous flow fields.

**FIG. 3.**
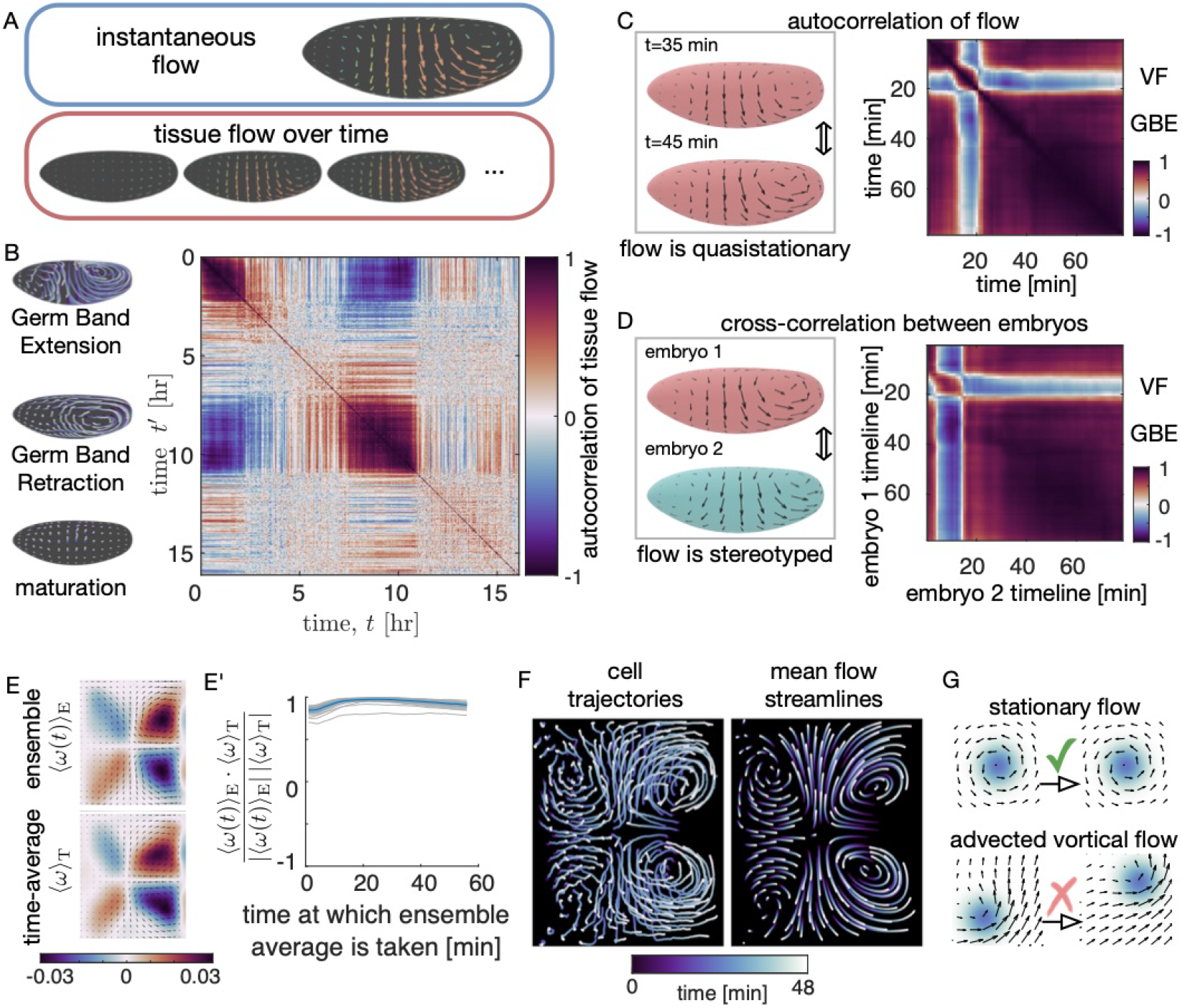
Tissue dynamics exhibit reproducible periods of quasi-stationary flow. *(A)* To characterize the characteristic periods of tissue flow, we compare the instantaneous tissue flow field against the timecourse of tissue flow during morphogenesis. *(B)* Comparing a single embryo’s flow against its timecourse of flow reveals blocks of quasi-stationary tissue flow during GBE and germ band retraction, here measured by the autocorrelation in vorticity. These blocks of high autocorrelation are punctuated by periods of low autocorrelation. *(C)* Zooming in on the autocorrelation at the onset of ventral furrow formation (VF) and GBE shows high correlation following VF. *(D)* Measuring the correlation in flow across embryos shows high cross-correlation in flow fields across samples. *(E)* The mean vorticity of each embryo during 56 minutes after GBE onset is strongly correlated with the ensemble-averaged vorticity across embryos for each developmental timepoint. The blue curve represents the average correlation between per-embryo and per-timepoint means across an ensemble of 57 embryos, and the blue shaded bar shows the standard deviation of these correlations across the ensemble. *(F)* Tracked trajectories in a *CAAXmCh* embryo shows a similar pattern of cell pathlines to the streamlines of the time-averaged and ensemble-averaged flow during GBE. *(G)* The quasi-stationary and highly stereotyped nature of the flow results from the nearly-stationary positions of the vortices in the embryo’s tissue flow. The positions of vortices are advected little compared to the motion of the tissue itself during GBE.

We developed a method to compare *integrated* flow patterns, cross correlating displacements of the tissue to one another using the same fast marching technique as in aligning live imaging of Runt stripes. In both cases, the morphology of the tissue marks its placement in the morphological timeline. We fix the reference timepoint of the integration as the time when GBE tissue flow starts to rise significantly (see Methods). As shown in Fig. 2F-H and the Supplementary Information, this approach leads to aligned tissue morphology.

Note that we use total tissue deformation, and not the instantaneous flow field, as a developmental land mark. In many contexts, instantaneous flow fields can be relatively constant in time (e.g. cells migrating with fixed speed), making them unsuitable as timestamps.

## III. CROSS-CORRELATION OF GLOBAL TISSUE DYNAMICS REVEALS PERIODS OF QUASI-STATIONARY FLOW PATTERNS

Expressing tissue motion as a time-dependent velocity field defined over the surface of the embryo allows us to compare tissue motion at different times or in different embryos. For example, we can directly compute the difference of two velocity fields.

We measured tissue flow in all live datasets included in the atlas using particle image velocimetry [21]. When we analyze these tissue flows, we find discrete periods of time in which the global pattern of tissue velocity remains remarkably stationary (Fig. 3B). We stress that the tissue itself is not stationary, but instead the *pattern of motion* is stationary: although the cells are moving across the embryo, the pattern generated by the flow is stationary during certain discrete stages of development.

To make this observation quantitative, the simplest step is to compare the dynamic velocity field to itself: comparing the velocity field at time *t* to the velocity field at a later time *t’* defines the autocorrelation function of the tissue flow. The magnitude of autocorrelation – measured using either the direction of motion or the rotational component of the flow – is high during discrete blocks of morphogenetic time, which correspond to established developmental episodes (Fig. 3B and Supplementary Information).

In particular, we see that the pattern of tissue flow changes little throughout germ band extension (GBE). The pattern of the autocorrelation matrix shows the quasi-stationary nature of the tissue motion pattern, shown in detail for a representative single embryo in Fig. 3C. Extending this approach to investigate cross correlations between embryos, we find remarkable similarity in the pattern of tissue flow between different samples Fig. 3D. This means that tissue flows during germ band extension are both quasi-stationary and stereotyped across multiple embryos.

To highlight a consequence of the high degree of self-similarity and reproducibility, we compared the ensemble-averaged rotational flow 〈*ω*(*t*)〉_*E*_ (i.e. average over different samples at the same time) to the time averaged vortical flow, 〈*ω*〉_*T*_, for an ensemble of 57 wild-type embryos. We observe very high correlation for nearly all of GBE (Fig. 3E-E’). This type of calculation, using a large number of samples aligned along a common timeline to study variations in time and across samples, is a typical use case of the dynamic atlas technology.

To validate our PIV-based approach, we tracked 6000 cells in an embryo with a cell membrane marker (CAAX-mCherry) and derived the kinematics from single-cell trajectories. As shown in the Supplementary Information, the kinematics based on cell-trajectories differ little from the PIV measures, and furthermore, they produce similar results in the cross-correlation between time-averaged and ensemble-averaged flow shown in Fig. 3E‘.

As a result of the quasi-stationary character of the tissue flow, there is a striking resemblance between the pattern of cell trajectories drawn in Fig. 3F, and the pattern of streamlines in a purely stationary, ensemble-averaged and time-averaged flow field. This result demonstrates that, despite massive movement of material, the morphogenetic program precisely coordinates a fixed pattern of tissue motion during discrete periods of morphological time.

## IV. KINEMATICS OF GBE FOLLOW SIMPLE TEMPERATURE-DEPENDENT SCALING

The rate of development is sensitive to environmental conditions, such as temperature. If morphological timing is perturbed, will the pattern of flow change only in its magnitude, or also in its orientation and sequence? Notably, Drosophila embryos can tolerate variations in temperature, allowing us to apply significant experimental perturbations.

To tune the rate of development, we measured the pattern of flow during GBE at a series of temperatures (17°C, 22°C, 27°C, with embryos viable at all temperatures). As shown in Fig. 4A, embryos cultured at 17°C demonstrated reduced tissue flow rate, while embryos at 27^°^ progressed through GBE more rapidly. Embryos cultured at 22^°^ adopted an intermediate flow rate. We find that quasi-stationary kinematics appear in each temperature condition. Moreover, temperature does not affect the spatial *pattern* of tissue motion encoded by the kinematics: the flow fields differ only in their magnitude.

**FIG. 4.**
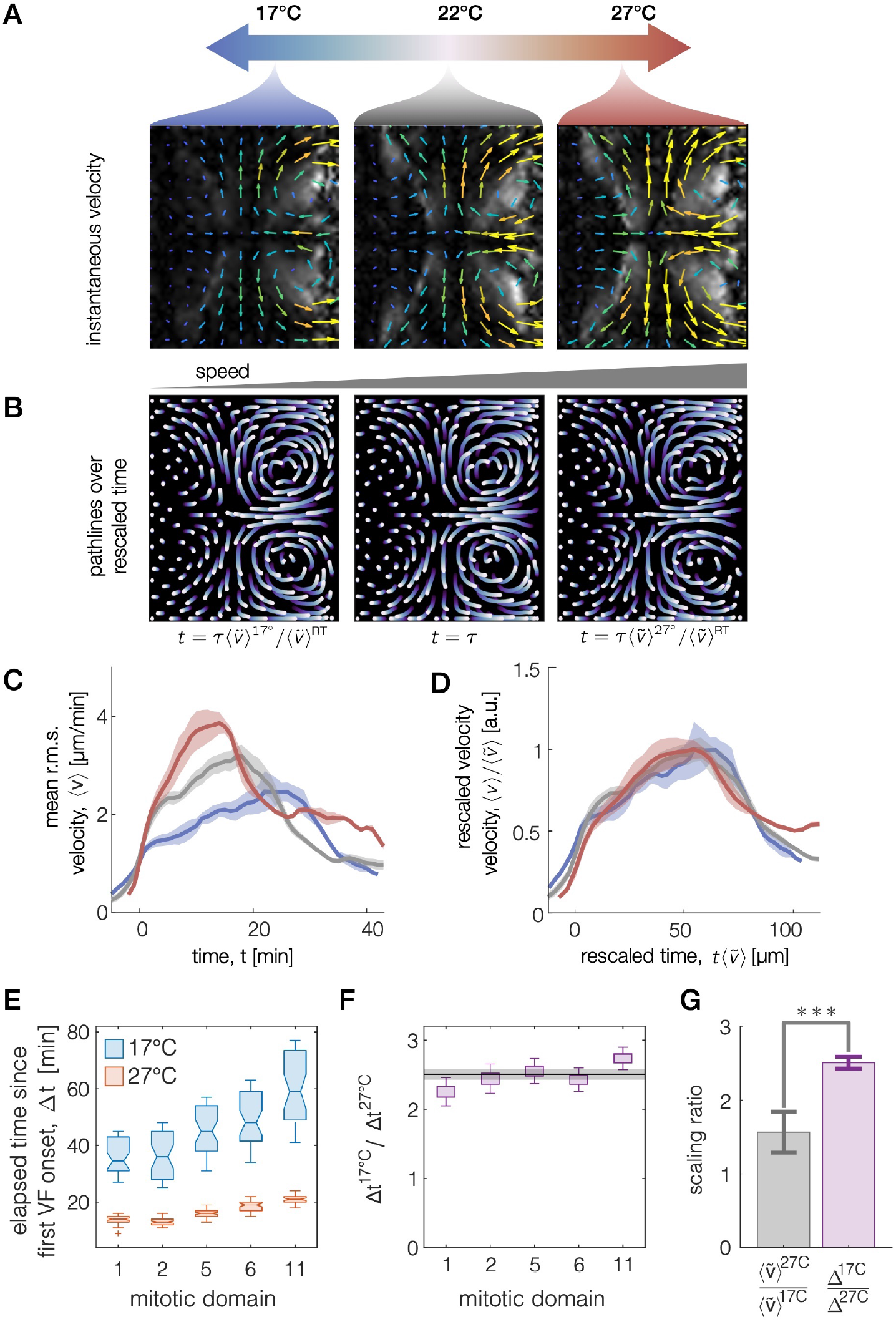
Kinematics of tissue flow scales with temperature without changing the pattern of deformation, but timing of mitotic domains scales differently with temperature. (*A*) The pattern of tissue velocity at different temperatures match in orientation and morphology, but differ in overall speed. Grey color represents flow vorticity. *(B)* Integrating velocities to follow tissue parcels over a timescale that is set by the overall speed results in nearly identical tissue pathlines. *t*= 0 corresponds to the onset of GBE tissue flow. Here, we use the maximum of the speed averaged over the embryo surface, 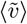, as the scaling factor, and *τ* is chosen to be 25 minutes at room temperature. *(C)* Measuring the average velocity across the surface of the embryo over time 〈*υ*〉 shows strong temperature dependence for embryos at 17°(blue), room temperature (gray), and 27°(red). (D) Rescaling time by the maximum velocity 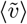 collapses all curves. This rescaling adjusts the velocities (rate of motion) by 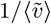 and stretches the time axis that parameterizes the velocities by 〈*υ*〉. (E) The time at which mitosis appears in different mitotic domains also shows strong temperature dependence. Here, the first 20 mitotic event timestamps are reported in bar-and-whisker plots for five different domains at reduced and elevated temperatures. *(F)*The time delay between the onset of ventral furrow formation and the onset of mitosis events shows a different scales with temperature by a factor of 2.5 ±0.1. (G) This temperature scaling ratio for mitosis is significantly different than the ratio that rescales tissue velocities. These measurements are largely independent since the mitoses contribute little to the tissue velocity during these stages (*** denotes *p* = 0.0005).

We hypothesized that the effect of temperature is simply to linearly accelerate or decelerate developmental time, much like playing an identical video at different speeds. This is consistent with the idea that the GBE program encodes target tissue deformations as a morphogenetic ‘checkpoint’, rather than prescribing, for example, a fixed time duration during which cells undergo rearrangement. The duration of tissue motions would need to vary in inverse proportion to the overall speed, to ensure that the final tissue deformation is conserved across conditions, and that the paths traversed by tissue patches are the same for all conditions.

To test this hypothesis, we integrated tissue motion for a duration set by the relative speed of the tissue flow, finding that the final pathlines are uniform across temperatures (Fig. 4B). As shown in the Supplementary Information, variations in tissue deformation across temperature conditions is not significantly different from variations within each condition. This leads to a parameter-free scaling, in which the embryo achieves the same final deformations (morphodynamic milestones) at variable rates, and all velocity curves collapse upon rescaling time by the maximum speed: *t* → max(〈*υ*〉) * *t* (Fig. 4C-D).

Does the same simple scaling govern all aspects of the morphogenetic program, or just tissue flows? Recent studies investigating temperature-dependent scaling have pointed to universal scaling across a survey of developmental processes [22, 23]. In contrast, other studies have found non-uniform temperature scaling for timing of the cell cycle and other subcellular processes, identifying divergent scaling rules for different components of the morphogenetic program [24, 25]. Motivated by these variations in scaling for mitosis, we investigated this question at the tissue scale by measuring the onset of mitosis in different domains across the embryo surface during GBE. These ‘mitotic domains’ are highly stereotyped in both relative timing and position [26].

To measure the rate of the ‘mitotic clock’ at each temperature, we calculated the time elapsed between the onset of division in different mitotic domains (see Supplementary Information). Remarkably, we find that the ratio of the time differences at different temperatures diverged significantly from the parameter-free scaling of the tissue motion. In particular, Fig. 4E-G shows that the relative timing of mitotic events across conditions varied by a factor of 2.5 ±0.1. In contrast, tissue flow velocities varied only by a factor of 1.5 ±0.3. The discrepancy is robust across different methods of measuring the time difference, such as choosing a different reference time (see Supplementary Information). This result opens further questions: why do cell cycle and tissue deformation timings scale differently, and how are these differences accommodated to produce viable embryos over a large range of temperatures?

## V. MORPHODYNAMIC ALIGNMENT OF A SHAPE-CHANGING VISCERAL ORGAN REVEALS SEQUENTIAL MORPHODYNAMIC MODULES IN COVARIANT MEASURES OF TISSUE DEFORMATION

Morphogenesis involves tissue dynamics not only on fixed surfaces such as the ellipsoidal surface of the early embryo, but also on more complex tissue geometries deforming in 3D space. At a later stage of embryogenesis, tissues that invaginate during gastrulation and in head involution come together to form the digestive tract [27]. During stages 15-16 of development, the midgut forms three constrictions in a sequence that divides the organ into four chambers, as shown in Fig. 5A [28–30]. This process is known to be stereotyped across embryos, but the quantification of its dynamics has until recently proven elusive, due to the complex shapes and inherently 3D deformations [30, 31]. Variability of the morphogenetic program between embryos remains largely uncharacterized [32].

**FIG. 5.**
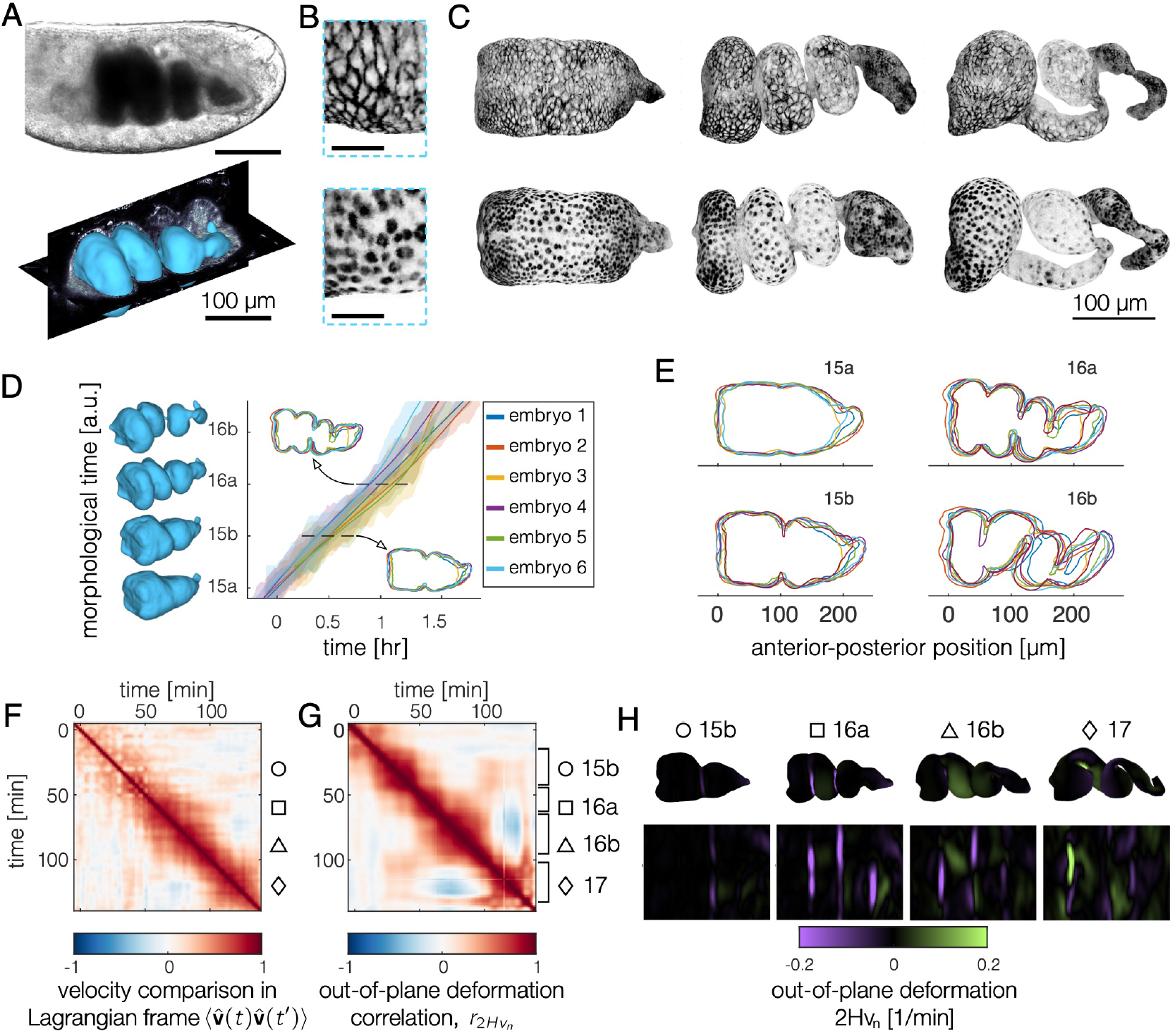
Morphodynamic atlas of the midgut via spatiotemporal registration reveals embryo-to-embryo shape variations and morphodynamic modules appearing in covariant measures of tissue deformation. *(A)* The midgut envelops to the embryo’s yolk and folds into chambers ~ 13-15 hours post fertilization. Extracting the endodermal (inner) surface of the gut using TubULAR [31] yields collections of dynamic organ shapes. Scale bars are 100 μm. *(B-C)* Our approach allows comparison across different classes of data, highlighted by plasma membrane marker and nuclear markers in the endodermal tissue surfaces rendered using TubULAR. Scalebar is 20 microns in (B). *(D)* Registration of one embryo’s midgut surface sequence to another allows pair-wise comparison and mapping of each embryo to a common morphological timeline. The embryos in our ensemble exhibited sterotyped morphogenesis with ~10% variation in the rate of development through morphological time. *(E)* Optimal spatiotemporal alignment allows comparison of embryo shapes across the ensemble. As morphogenesis proceeds, midguts become increasingly unique in their morphology. *(F)* Autocorrelation of the 3D tissue velocity for a representative embryo shows little temporal structure, leaving a diffuse streak of correlation along the diagonal. *(G)* In contrast, for measurements in the tissue frame of reference, block-like structures emerge in sequence along the diagonal, here for the auto-correlation of the out-of-plane deformation. *(H)* Snapshots of the covariant out-of-plane deformation for each morphodynamic module shows stereotyped deformation for each morphogenetic stage.

**FIG. 6.**
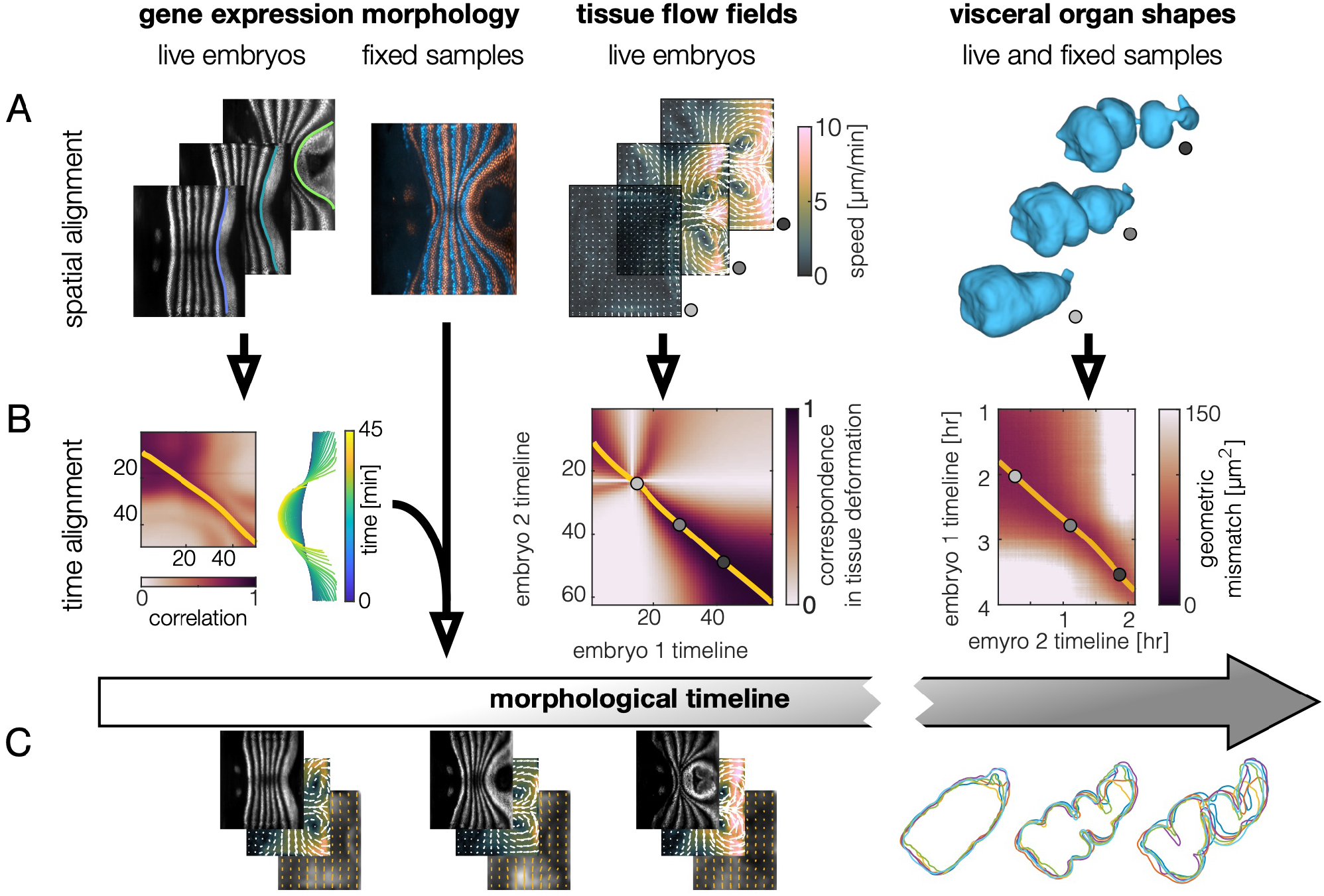
Comparing morphodynamic features across datasets generates a morphodynamic atlas. *(A)* Extracting geometric deformations and *(B*) cross-correlating these features between samples defines *(C)* a consensus morphological timeline. Live expression profiles define a consensus morphological timeline and temporal sequence of Runt stripe geometries. Immunostaining against Runt in all fixed samples within the atlas defines their morphological time. Comparing tissue flow fields in dynamic datasets without visible Runt expression similarly define correspondences which we integrate into the common morphological timeline. Sequences of midgut shapes similarly define a 3D geometric indicator of morphological time.

During this stage of development, the tissue at the embryo surface shows very little coherent motion compared to GBE, and the autocorrelation of the tissue flow on the ectodermal surface from Fig. 3B during this time is nearly zero. Will characterization of the complex tissue motion deep inside the embryo reveal morphodynamic modules?

To test this, we here extend our spatiotemporal registration and examination of velocity correlations during morphogenesis to the development of complex shapes in the midgut. We use level sets approaches of the TubULAR package [31, 33] to extract a surface that penetrates ~2.5 μm within the apical surface of the endoderm, along the surface that intersects endodermal nuclei. In contrast to the case of germ-band extension, these surfaces are highly dynamic, demanding additional steps for spatial registration.

With sequences of each embryo’s midgut surface in hand, we find the closest match between each pair of surfaces using iterative closest point registration (see Methods). This algorithm finds the combination of rotation and translation that best maps two 3D surfaces onto each other. This morphological approach allows alignment across embryos with different fluorescently tagged proteins (Fig. 5B).

With spatial registration performed, we can use a quantitative comparison of organ shape to temporally align the process of midgut morphogenesis across embryos. Performing pairwise alignment across embryos leads to a consensus timeline of morphology shown in Fig. 5C. For our ensemble of embryos, the rate of development through morphological stages varied by ~ 10%. Fig. 5D shows cross-sections of these embryos in the lateral plane during four stages of development. The match between midgut shapes becomes less stereotyped as morphogenesis proceeds, as evident in quantification of the shape variation across our ensemble (Fig. 5D and Sup-plementary Information).

We then turned to whether the tissue kinematics exhibit periods of quasi-stationary motion like in the early embryo’s axis elongation. Fig. 5E shows that unlike the earlier velocity autocorrelation maps of GBE, here the full 3D velocity’s autocorrelation shows relatively little structure: while dynamics vary continuously and thus show some degree of correlation with similar timepoints, no distinct morphodynamic modules are present. This makes intuitive sense, as coordinate directions in the tissue frame of reference rotate as the surface deforms in 3D space.

We find more structure, however, in the autocorrelation of measures of tissue deformation computed in the tissue’s frame of reference. Covariant measures of deformation – which make reference to the tissue’s orientation in 3D space – do show distinct morphodynamic modules (Fig. 5F). While the *time-averaged* autocorrelation of both classes of measurements display similarly prolonged correlation timescales of ~25 minutes (see Supplementary Information), moving into the reference frame of the organ to evaluate covariant measures of tissue motion highlights boundaries between modules which are hidden in the full 3D velocity field measurement. Within these modules, the autocorrelation forms box-like shapes of high correlation arranged along the diagonal in Fig. 5F.

Examination of the out-of-plane deformation during these modules shows qualitatively distinct programs, illustrated for a representative embryo in Fig. 5G. During the first module, the middle constriction forms (purple streak bisecting the organ). In [30], this module is denoted as stage 15b. Subsequently, out-of-plane deformation appears at the anterior and posterior constriction locations. This module shares ~50% correlation with the stage 15b, as the middle constriction continues to deepen. In [30], this second module is denoted as stage 16a. A third module, which shares significant correlation with 16a but not with 15b, then follows, marking the absence of significant divergence or out-of-plane deformation near the posterior constriction and a breaking of left-right symmetry in the second chamber. Finally, constrictions are complete, and a new pattern of deformation emerges during stage 17, in which the organ continues to coil into a helical configuration. Similar modules arise upon examination of other covariant measures of tissue deformation (see Methods).

This analysis demonstrates that while the morphogenetic program is encoded in stepwise modules, not all aspects of tissue velocity will exhibit these quasistationary features. More generally, our approach offers a systematic strategy for discovering mechanisms for encoding morphogenetic processes.

## VI. DISCUSSION

We have constructed an open-source morphodynamic atlas of Drosophila development – which integrates dynamic and static datasets capturing patterning gene expression, cytoskeletal patterning, and nuclear and membrane markers – and an associated computational platform for querying and interfacing with the atlas. Using automated extraction of measures of tissue geometry, we align all datasets to a common morphological timeline covering GBE and midgut morphogenesis. Our fast-marching based timeline creation algorithm automatically takes variations in developmental speed into account. This resource provides a testbed for rapid hypothesis testing and quantitative modeling. In addition, our approach to indexing by morphology and cross-correlating data across samples is valuable as a tool for addressing variability within and between ensembles.

Comparing *in toto* tissue velocities across time and across embryos shows stereotyped modules of quasistationary tissue deformation both during body axis elongation and midgut morphogenesis. Temperature perturbations revealed that the duration of the quasi-stationary flow pattern during body axis elongation follows a simple, parameter-free scaling in which the same final tissue deformation is achieved at a temperature-dependent rate.

This scaling suggests that the morphogenetic program encodes total tissue deformation, i.e. that the duration for which the morphogenetic modules revealed by our correlation analyses are active is determined by the accumulated tissue deformation. Mitotic events, however, followed a distinct scaling with temperature, raising the question of whether the two are synchronized at a subsequent morphogenetic checkpoint. Such checkpoints could occur at “pauses” between morphogenetic blocks. Differences in scaling for different aspects of morphogenesis have been noted before in other contexts [34].

The reproducibility of tissue flow is reminiscent of the well-documented spatial precision of gene expression patterns [35, 36]. These earlier studies showed gene expression levels to be spatially patterned with single-cell precision. Tissue kinematics are likewise strongly correlated in a fixed, geometric (Eulerian) reference frame, despite massive motion of the material. This result is in line with the recent finding that the anisotropic pattern of myosin appears to be controlled by static, non-deforming cues during GBE tissue motion [19].

## VII. ACKNOWLEDGEMENTS

We thank Eric Wieschaus, Fridtjof Brauns, and Dillon Cislo for insightful discussions. Eric Wieschaus contributed stocks and reagents. This work was funded by NIH Grant No. R35 GM138203, and in part by the National Science Foundation under Grant No. NSF PHY-1748958. NPM acknowledges support from the Helen Hay Whitney Foundation.

